# Extraversion is associated with lower brain amyloid deposition in cognitively normal older adults

**DOI:** 10.1101/2022.01.23.477394

**Authors:** Hwamee Oh

## Abstract

Emerging evidence suggests that some personality traits may link to the vulnerability to or protection for Alzheimer’s disease (AD). A causal mechanism underlying this relationship, however, remains largely unknown. Using ^18^F-Florbetaben positron emission tomography (PET) binding to beta-amyloid (Aβ) plaques, a pathological feature of AD, and functional magnetic resonance imaging (fMRI), we investigated pathological and functional correlates of extraversion and neuroticism in a group of healthy young and older subjects. We quantified a level of brain Aβ deposition in older individuals. Brain activity was measured in young adults using a task-switching fMRI paradigm. When we correlated personality scores of extraversion and neuroticism with these pathological and functional measures, higher extraversion, but not neuroticism, was significantly associated with lower global Aβ measures among older adults, accounting for age and sex. This association was present across widespread brain regions. Among young subjects, higher extraversion was associated with lower activity during task switching in anterior cingulate cortex, left anterior insular cortex, left putamen, and middle frontal gyrus bilaterally, while higher neuroticism was associated with increased activity throughout the brain. The present results suggest that possibly via efficient neuronal activity, extraversion, one of lifelong personality traits, may confer the protective mechanism against the development of Aβ pathology during aging.

## Introduction

Alzheimer’s disease (AD) is characterized by the pathological accumulation of beta-amyloid (Aβ) peptides and tau-protein neurofibrillary tangles, followed by neuronal injury and severe cognitive impairment^1, 2^. Neuropathological and epidemiological studies suggest that lifestyle factors such as education, social network, diet, and cognitive and physical activities, may confer an increased risk or conversely a protective mechanism for the clinical manifestation of AD as well as the development of AD pathology^3–5^. The observed association between lifestyle factors and clinical and pathological manifestation of AD points to the possibility that the vulnerability to AD pathology and onset of AD symptoms may develop over the lifespan. Although relatively less attention has been given compared with these lifestyle factors, emerging evidence suggests that an individual’s personality, a trait that is found to be relatively stable across the lifespan, relates to the increased risk or protection of AD^6–9^. A neuro-mechanistic explanation linking the personality traits to AD, however, is largely unknown. Personality often interacts with the proposed lifestyle factors such as cognitive and physical activity and social network^3,10^. Therefore, the understanding of the neural mechanism linking personality to AD risk will be an important step in disentangling specific effects of complex lifestyle variables that may act simultaneously across the lifespan in the development of resilience or vulnerability to AD.

Personality traits are an individual’s styles and tendencies of interacting with external events. There is general consensus that adult personality can be defined by five factors, known as the Big Five: neuroticism, extraversion, openness to experience, agreeableness, and conscientiousness^11^. Among personality traits, neuroticism has mostly linked to an increased risk of dementia, while conscientiousness have been related to healthier lifestyle choices^12,13^. Extraversion and openness have been associated positively with intellectual ability which in turn moderates the severity of clinical symptoms^10,14,15^. Over the course of AD, the level of neuroticism and extraversion has changed, showing higher level of neuroticism and lower level of extraversion in AD patients compared to premorbid status. Collectively, personality traits and AD risk have been linked; however, what neural mechanism underlies this relationship remains to be elucidated.

In the process of AD, Aβ accumulation, a pathological feature of AD, is considered as a relatively early event that antedates more than a decade the appearance of clinical symptoms^2,16^. Findings from both animal and human studies suggest that Aβ accumulation relates to neural activity and physiology early in life before plaque accumulation^17–20^. For example, regional concentration of interstitial fluid Aβ in young mice was associated with lactate production indicating neuronal activity and was further related to amyloid-β plaque deposition in the same brain regions in APP transgenic aged mice^18^. In humans, Aβ accumulation in older adults topographically overlaps with brain regions showing higher metabolic rates in young adults^19,20^. Interestingly, neuroimaging studies examining personality trait-based phenotypes have shown differential brain activity patterns in relation to personality types. Among personality traits, introversion/extraversion and neuroticism have been most influential and frequently studied dimensions of human personality. Across a range of emotional and cognitive tasks, neuroticism has been associated with increased brain activity in the medial frontoparietal network including medial prefrontal cortex and precuneus^21–23^. On the other hand, extraversion has been associated with decreased activity across the brain reflecting neural processing efficiency or reduced self-consciousness^23,24^. Considering the epidemiological, neuropathological, and neurocognitive findings, personality traits, possibly via differential regulation of regional neural activity across the lifespan, may contribute to an individual’s risk to the buildup of AD pathology during aging.

Based on the literature linking across personality traits, neural activity, and Aβ deposition, we sought to directly test the relationship between personality and Aβ pathology and a potential neural mechanism underlying this relationship. Specifically, we tested whether extraversion is associated with lower Aβ pathology measured by^18^F-Florbetaben PET in cognitively normal older adults and reduced neural activity in the medial frontoparietal cortex during executive control in young adults. To confirm the specificity of the effect of extraversion on Aβ deposition via neural activity, we also examined the relationship between neuroticism and brain activation pattern in young adults. We hypothesized that cognitively normal older adults with higher extraversion scores would show lower Aβ pathology and that higher extraversion in young adults would be associated with reduced activity in the medial frontoparietal cortices during executive control. With neuroticism, we hypothesized that higher neuroticism scores in young adults would be associated with increased activity in the medial frontoparietal network. Because Aβ deposition is highly regulated by neural activity, a differential pattern of brain activity in association with extraversion and neuroticism in young adulthood would provide a link between personality and Aβ pathology, which may form across the lifespan and would be observed among older adults.

## Materials and Methods

### Participants and neuropsychological assessment

Thirty-three healthy young (age range:20-30) and 57 cognitively normal older adults (age range:60-70) participated in the study. Demographic details are provided in Table 1. Participants were a subset of subjects who participated in our previous study^25^ and whose personality test measures were available. Detailed description of subject characteristics can be found in our previous report^25^. Briefly, all subjects were recruited via a market mailing procedure and older subjects were classified as either “Amyloid-positive” (Aβ+O) or “Amyloid-negative” (Aβ-O) based on amyloid PET measurements as described below. All subjects provided informed consent in accordance with the Institutional Review Boards of the College of Physicians and Surgeons of Columbia University. Participants were paid for their participation in the study.

**Table 1.** Subject characteristics.

|  | **Young** | **Amyloid– Old** | **Amyloid+ Old** |
| --- | --- | --- | --- |
| N | 33 | 43 | 14 |
| Age | 26.4 (3.1) | 65.3 (2.9) | 65.4 (3.7) |
| Edu | 15.5 (1.9) | 16.4 (2.5) | 15.9 (1.8) |
| Sex(F,%) | 23 (69%) | 21 (49%) | 8 (57%) |
| Amyloid index |  | 1.16(0.05) | 1.36(0.1) |
| Extraversion | 3.40 (0.8) | 3.34(0.7) | 2.96(0.8) |
| Neuroticism* | 2.90(1.0) | 2.57(0.7) | 2.17(0.8) |
| Memory** | 0.5 (0.4) | -0.4 (0.6) | -0.3 (0.8) |
| Processing Speed** | 0.8 (0.6) | -0.4 (0.7) | -0.3 (0.6) |

Values (SD). \* $p < 0.05$ , \*\* $p < 0.01$

A comprehensive battery of neuropsychological tests was administered to all participants to assess cognition. Among cognitive domains, we computed a composite score for processing speed/attention using Wechsler Adult Intelligence Scale-Third Edition (WAIS-III) Digit Symbol subtest^26^, Trail Making Test Part A^27^(inversed value), and Stroop Color Naming test-Color naming in 90 sec^28^ and a composite score of episodic memory combining scores from Selective Reminding Test (SRT)^29^-long term storage, SRT-continued recall, and SRT-recall at the last trial.

For extraversion and neuroticism scores, we used a subset of the international personality item pool (IPIP) Big-Five personality inventory^30^, which additionally consists of items for openness, agreeableness, and conscientiousness. The IPIP inventory is a 50-item questionnaire, with 10 items for each personality type, in a 5 point Likert scale (1=“Very Inaccurate”, 2=“Moderately Inaccurate”, 3=“Neither Inaccurate nor Accurate”, 4=“Moderately Accurate”, 5=“Very Accurate”). An averaged score for each personality type represented each subject’s personality score.

As summarized in Table 1, older subjects were more educated, although at a trend level, than young subjects *(T* (88)=-1.8, *p*=0.07). Aβ+O vs. Aβ-O groups did not differ in age, gender, education, dementia rating scale (DRS), or IQ estimated by AMNART. With cognitive composite scores, young subjects showed significantly higher scores in memory than Aβ-O subjects, while no other group differences were found for both memory and processing speed composite scores. For extraversion and neuroticism scores, young subjects showed significantly higher scores in neuroticism compared to Aβ+O subjects. Personality scores for five factor personality domains are summarized for all subjects and by age group in Supplementary Table 1.

### ^18^F-Florbetaben-PET acquisition, image processing and analysis

^18^F-Florbetaben was donated by Piramal (Piramal Pharma Inc.). Detailed information on^18^F-Florbetaben PET data acquisition and image processing can be found in our previous report^25^. Briefly, scans were performed on a Siemens MCT PET/CT scanner in dynamic, three-dimensional acquisition mode over 20 minutes (4X5 min frames) beginning 50 min following the bolus injection of 10 mCi of^18^F-Florbetaben. Dynamic PET frames (4 scans) were aligned and averaged to form a mean PET image that was coregistered to each individual’s structural T1 image in FreeSurfer space. The standardized uptake value ratio (SUVR) was calculated for selected cortical regions encompassing frontal, temporal, parietal, and anterior/posterior cingulate cortices with cerebellar gray matter as a reference region^31^,^32^. Mean SUVR values from these lobar ROIs constituted a global amyloid index for each subject. Amyloid positivity of elderly subjects was determined using a K-means clustering method as previously reported^33^. Fourteen and 43 older subjects were classified as Aβ+ and Aβ-subjects, respectively. The proportion of Aβ+ subjects in our sample is comparable to the reports from other studies^34^.

In order to assess the relationship between personality trait and Aβ deposition, we performed 2 analyses: (1) multiple regression model using a mean cortical Aβ measure as a dependent variable and personality scores as an independent variable and (2) the wholebrain voxel-wise analysis using general linear model treating voxel-wise Aβ level as a dependent measure and personality scores as an independent measure. The whole brain family-wise error was cluster corrected to *p*<0.05 (two-sided) using a cluster forming threshold of *p*<0.05. Age and sex were controlled in all analyses.

### fMRI experimental task

To assess a brain activation pattern in association with extraversion and neuroticism, we used the executive contextual task adapted from Koechlin et al (2003) which is described in detail in our previous report^25^. Briefly, the fMRI task was designed to assess executive control function in a block-design in which either single or dual task condition was assigned (Supplementary Figure 1). During fMRI scans, a letter in either red or green appeared on the screen and subjects were asked to do vowel/consonant judgment for a green letter and lower/upper-case judgment for a red letter. For a single task condition, either green or red letter appeared throughout a block so that subjects had to do only one type of task throughout the task block. For a dual task condition, color of letters changed between green and red so that subjects had to switch between two tasks within a task block accordingly. Twelve letters were presented for a maximum of 2400 msec each within a block that was 33.5 sec in duration. A one third of letters in a block appeared in white, for which no response was required. Each functional run consisted of four task (2 single-task and 2 dual-task) blocks and 2 resting condition blocks during which no stimuli were presented and no response was required; in total, the scan session was composed of 6 functional runs. The executive contextual task fMRI session lasted approximately 26 min.

Response time (RT) of correct trials and proportion correct for single and dual task conditions were calculated for each individual and reported as behavioral measures.

### MRI data acquisition

Participants underwent MRI using a 3T Philips Achieva System equipped with a standard quadrature headcoil. High-resolution T1-weighted magnetization-prepared rapid gradient echo (MPPAGE) scans were collected axially for each subject (TR=6.6 ms, TE=3 ms, flip angle=8°, field of view (FOV)=256X256 mm, matrix size: 256X256 mm, slices: 165, voxel size=1X1X1 mm^3^). For the executive context task scans, 111 volumes of functional images were acquired in each run (6 runs in total) using a T2*-weighted gradient-echo echo planar images (EPI) sequence (TR=2000 ms; TE=20 ms; flip angle=72°; FOV=224X224 mm; voxel size=2X2 mm; slice thickness=3mm; duration=3.5 min). Each functional volume consisted of 41 transverse slices per volume. Four dummy volumes acquired at the beginning of each functional run were discarded from the data set before image processing and analysis.

### Structural MRI image processing

For each subject, a single structural T1 image was processed through FreeSurfer v5.1 to implement region of interest (ROI) labeling following the FreeSurfer processing pipeline (http://surfer.nmr.mgh.harvard.edu/). Briefly, structural images were bias field corrected, intensity normalized, and skull stripped using a watershed algorithm, followed by a tissue-based segmentation, defining gray/white matter and pial surfaces, and topology correction^35^. Subcortical and cortical ROIs spanning the entire brain were defined in each subject’s native space^36^.

### fMRI image processing and analysis

All fMRI analyses were performed with SPM8 (Wellcome Departmentof Imaging Neuroscience, London, UK). Detailed description of fMRI data preprocessing and subject-level analysis can be found elsewhere^25^. To identify brain regions that show task-related activity associated with extraversion and neuroticism in young adults, we focused on task-related activity from three comparison/contrasts: single-task condition compared with baseline, dual-task condition compared with baseline, dual vs. single task contrast. Using general linear models, each comparison/contrast-related activity was regressed on extraversion and neuroticism scores, with gender as covariates of no interest. The whole brain family-wise error was cluster corrected to *p*<0.05 (two-sided) using a cluster forming threshold of *p*<0.05.

### Non-image data analysis

All non-image data anlaysis was conducted using SPSS v.22. The relationship between personality trait and Aβ deposition was assessed using multiple regression with a mean cortical Aβ measure as a dependent measure and personality scores as an independent measure. The direct correlation across all older adults was further probed using analysis of covariance (ANCOVA) to assess an interaction effect of amyloid positivity status and personality scores on global Aβ measures. The association between personality scores and cognition was assessed using cognitive composite scores of memory and processing speed as well as executive control task performance (e.g., RT and proportion accuracy). Age and sex were controlled as covariates of no interest.

## Results

### Extraversion is associated with lower amyloid deposition among cognitively normal older adults

We assessed the relationship between personality trait and Aβ deposition using a mean cortical and whole-brain voxel-wise Aβ measures. Using multiple regression model with a mean cortical Aβ measure, we found that, among cognitively normal older adults, higher extraversion scores were significantly associated with lower global amyloid index (*β*=-0.31, *p*<0.05) (Fig. 1B). In addition to a global Aβ measure, we found the similar results using local amyloid deposition in posterior cingulate cortex (*β*=-0.37, *p*< 0.05) (Fig. 1C). Neuroticism scores, however, were not significantly associated with amyloid burden (*p*>0.1). A significant relationship between extraversion scores and amyloid burden was mainly driven by a significant association within Aβ+ group, as revealed by a significant interaction between extraversion scores and amyloid positivity status on global amyloid index (*F*=4.2, *p*<0.05). Next, we performed the whole-brain voxel-wise analysis to further probe regional specificity for Aβ deposition in relation to extraversion scores. The cortical regions wherein the level of Aβ deposition significantly related to extraversion score are shown in Fig. 1A (cluster *p*<0.05), indicating that higher extraversion scores are associated with lower Aβ deposition across frontal, parietal, and temporal cortices. These brain regions highly overlap with regions that typically accumulate more Aβ than other brain areas.

**Figure 1.**
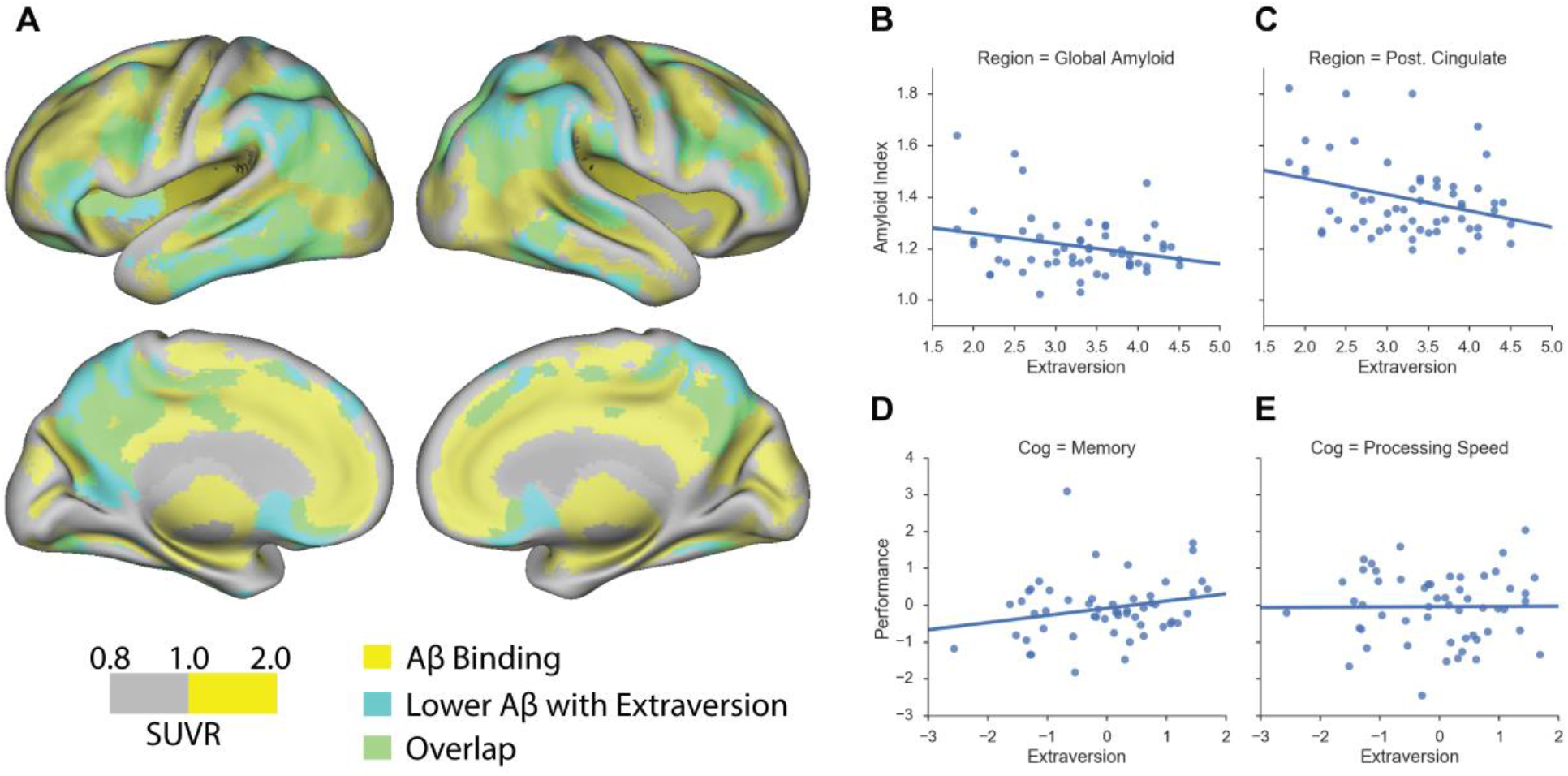
Extraversion is associated with lower amyloid deposition in cognitively normal older adults. A. Association between voxel-wise standardized uptake value ratio (SUVR) measures and extraversion among cognitively normal older adults. Scale represents voxel-wise SUVR measures. Regions highlighted in yellow represent voxels whose SUVR values equal to or are greater than 1. Regions highlighted in cyan represent suprathreshold voxels showing significant negative association between extraversion and amyloid burden. Regions highlighted in green indicate an overlap between amyloid deposition and a significant negative association with extraversion scores. B. Extraversion scores are associated with lower amyloid deposition as quantified by a global amyloid index that represents mean cortical amyloid burden. C. A significant negative association between extraversion and amyloid deposition is found using mean amyloid accumulation in posterior cingulate cortex (Post. Cingulate). D. Extraversion scores (residual values) are associated with better, although at a trend level, memory performance (residual values) among cognitively normal older adults. E. Extraversion scores are not associated with processing speed. Age and sex were controlled in the analyses.

Using the two approaches, we further performed exploratory analyses to test an association between global Aβ deposition and other personality scores: conscientiousness, openness, and agreeableness. No significant association was found between global Aβ level and these personality measures (*ps*>0.1). Age and sex were controlled in all analyses.

### Differential brain activity is associated with extraversion and neuroticism in young adults

For fMRI task behavioral measures, we examined an association between extraversion/neuroticism scores and response time and proportion correct during single and dual-task conditions. Neither extraversion nor neuroticism was related to any behavioral performance measure (*ps*>0.1).

For task-evoked brain activation, overall, higher extraversion scores in young adults were related to less brain activation in single task compared to baseline and dual task compared to baseline (Fig. 2A). Specifically, in the single task compared to baseline activity, higher extraversion was associated with reduced activation in anterior cingulate, medial frontal cortex, and lateral middle frontal cortex. In the dual task compared to baseline activity, a similar but more extended pattern of activity was found in association with extraversion scores. In the dual vs. single task contrast, there was no suprathreshold activity. With neuroticism, we found extensive increases in brain activity in association with higher neuroticism (Fig. 2B). Across the activation maps, higher neuroticism was associated with increased activity in anterior cingulate cortex, anterior frontal cortex, right insular cortex, and lateral inferior and middle frontal cortex, posterior cingulate cortex, right superior frontal cortex, caudate, and thalamus, while more extensive regions showed a positive association between neuroticism and brain activity for the dual task compared to baseline activity.

**Figure 2.**
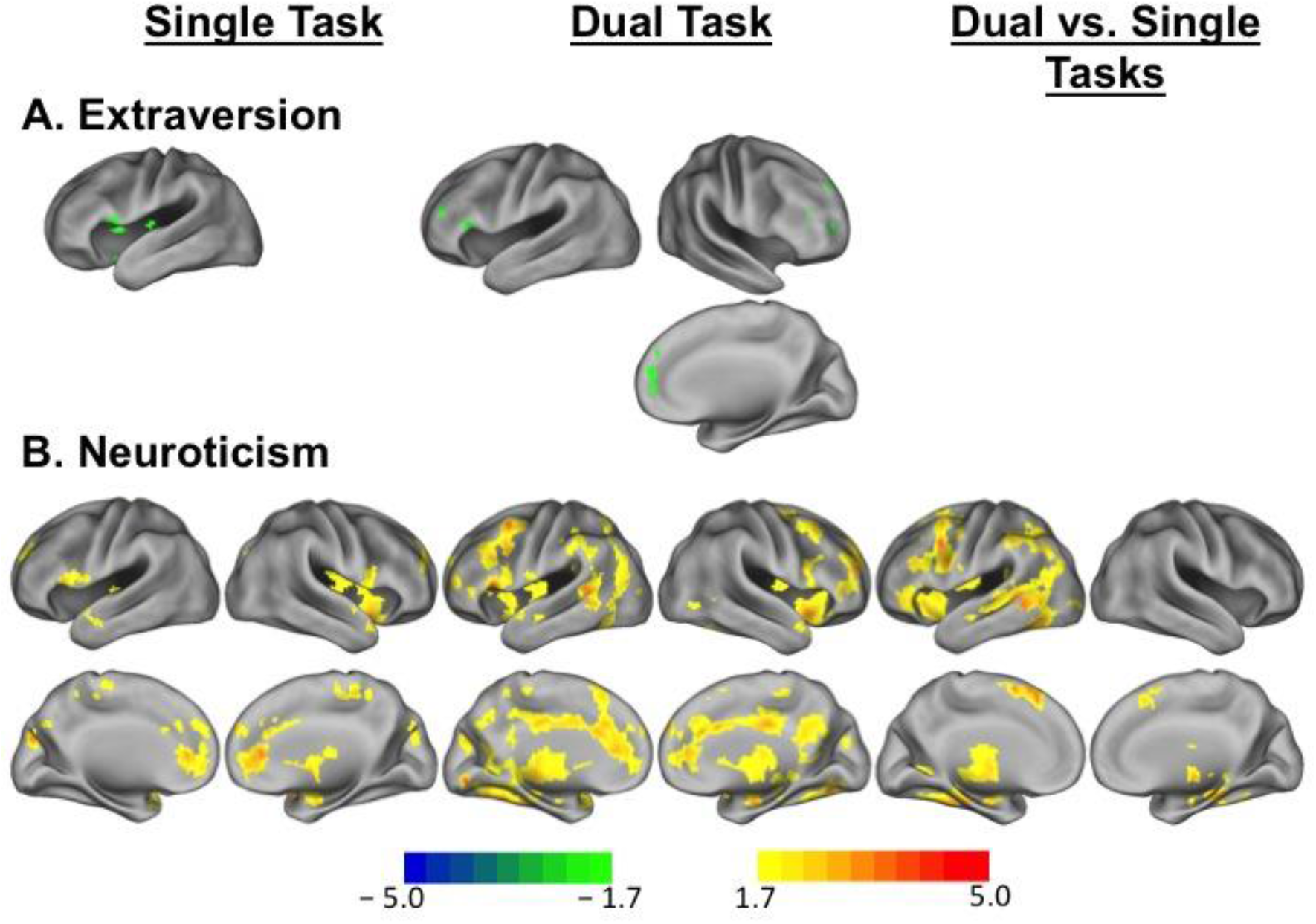
Differential brain activity patterns in association with extraversion and neuroticism among young adults. A. Task-evoked brain activity during executive control tasks in relation to extraversion scores among young adults. Regions in green indicate suprathreshold voxels showing lower brain activity in relation to higher extraversion scores for single task vs. baseline and dual task vs. baseline comparisons. B. Task-evoked brain activity during executive control tasks in relation to neuroticism scores among young adults. Regions in warm colors indicate suprathreshold voxels showing increased brain activity in relation to higher neuroticism scores for single task vs. baseline and dual task vs. baseline comparisons and dual vs. single task contrasts. Scale represents *T* values.

### Association between personality and cognitive performance

We conducted several exploratory analyses to better understand the demographic and psychosocial factors and cognition that relate to personality traits of extraversion and neuroticism in healthy young and cognitively normal older adults. Among cognitively normal older adults, we found no association between extraversion and neuroticism, controlling for age and sex. Extraversion was not correlated with age, but significantly correlated with sex, such that women showed significantly higher extraversion compared to men (*β*=0.4, *p*=0.003). Neuroticism was not correlated with sex, but significantly correlated with age, showing that the older the age is, the lower the neuroticism scores are (*β*=-0.33, *p*=0.02). A significant relationship between extraversion and memory composite scores was found at a trend level (*β*=0.28, *p*=0.08; Fig. 1D), but no significant relationship was found between extraversion and processing speed composite score (*β*=0.03, *p*>0.1; Fig. 1E). Neuroticism was not related to any cognitive composite measure. In relation to other personality traits, extraversion was significantly associated with openness (*β*=0.3, *p*=0.02), controlling for age and sex. Neuroticism was significantly negatively associated with agreeableness (*β*=-0.27, *p*=0.046) as well as conscientiousness (*β*=-0.35, *p*=0.01), controlling for age and sex.

Among young subjects, there was no association between extraversion and neuroticism (*p*>0.1). Neither age nor sex was associated with extraversion or neuroticism (*ps*>0.1). With neuropsychological composite scores, personality measures were not associated with any neuropsychological score (*ps*>0.1). In relation to other personality traits, extraversion was significantly associated with agreeableness (*β*=0.5, *p*=0.01) as well as conscientiousness (*β*=0.4, *p*=0.035), controlling for age and sex. Neuroticism was not associated with any other personality trait scores (*ps*>0.1).

## Discussion

In the present study, we report a novel finding showing that higher extraversion scores are associated with lower Aβ deposition across the widespread brain regions among cognitively normal older adults. When we examined personality-related differences in task-evoked brain activity among young adults, we found a significant dissociation in brain activity as a function of personality traits: extraversion was related to lower brain activity, in particular, in the medial and lateral frontal cortex and anterior cingulate cortex, while neuroticism was associated with increased activity throughout the brain. Our results are consistent with a metabolism hypothesis of Aβ pathology^37^ and provide neural evidence linking personality traits to a source of risk and protective factors of AD that are formed throughout the lifespan.

### Lifestyle factors as reserve or risk of AD

There is great interest in understanding individual differences that influence the vulnerability of individuals to age-related neurodegenerative diseases such as AD. A large body of neuropathological and epidemiological studies suggests that lifestyle factors such as education, social network, diet, and cognitive and physical activities, may affect the clinical manifestation of AD as well as the development of AD pathology. The relationship between lifestyle factors and AD has been commonly investigated with at least three hypothesized pathways. The pathway that received the most attention is its modulatory pathway that acts on the connection between disease pathology and clinical symptoms, which has been termed as cognitive or neural reserve^3,4^. The other equally plausible pathways are to influence directly either to the buildup of AD pathology or to clinical symptoms independent of AD pathology^3,6,38–44^. Epidemiological and neuropathological data from large cohort studies are collectively more supportive of the modulatory effect of lifestyle factors on clinical manifestation of AD both in cognitively normal older adults^45^ and AD patients^3, 46^.

### Personality as a risk and/or protective factor for AD

Compared with the role of education and cognitive aspects of lifestyle factors in the occurrence of AD, less attention has been given to the role of personality and related social engagement factors in AD neuropathology and dementia status. The most common model of personality organizes personality traits into five factors: neuroticism, extraversion, openness to experience, agreeableness, and conscientiousness^11^. Studies have consistently found that measures of these personality traits predict AD risk and important health outcomes of their lives. Neuroticism, an individual’s increased tendency to negative affectivity, anxiety, and distress proneness, has been shown to predict higher psychological distress, cognitive decline, mortality, and increased risk of AD^12,47^. Loneliness, which is defined as perceived isolation, was robustly associated with cognitive decline and development of AD^8^. Personality-related risk and beneficial effects on the onset of clinical symptoms and AD pathology were further supported once a possibility of premorbid personality changes has been accounted for^6^. Using a quantitative measure at both global and voxel-wise level, our results provide evidence showing a significant association between amyloid deposition and extraversion among older adults who are yet clinically intact.

A central motivation of the present study was to elucidate the neural basis underlying the impact of personality traits on AD pathology. Some morphological studies have shown larger gray matter (GM) volume in association with low neuroticism and higher extraversion, openness, agreeableness, and conscientiousness mostly in higher-level association cortices^48^. Because these results remained identical across different time points, findings further suggest that distinct personality traits are associated with stable individual differences in GM volume and potentially act as a risk or protective factor by presenting neural reserve during brain aging.

A more direct link in relation to AD pathology can be drawn from brain activation studies showing that personality traits relate to unique patterns of brain activity in response to emotional or cognitive events or stimuli. Neuroticism is associated with negative mood states, sensitivity to negative information, negative appraisal and vulnerability to psychopathology, which commonly relate to the sustained processing of negative information. By assessing sustained patterns of activity within a brain region implicated in emotional self-evaluation and appraisal, such as the medial frontal cortex, studies have found that higher scores of neuroticism are associated with greater sustained patterns of brain activity in the medial frontal cortex when responding to blocks of negative facial expressions^49^. In the cognitive domain, controlled processing mechanisms have been linked to the dorsal anterior cingulate cortex (dACC) and the reactivity of dACC has been directly related to neuroticism, due to its role in discrepancy detection^23^. With regard to extraversion, studies consistently show lower baseline activity with higher extraversion and task-related decreases in the lateral and medial frontoparietal network, typically associated with task-focused controlled processing, reflecting neural-processing efficiency, and reduced self-consciousness^23,24,50^. Our findings of differential activity as a function of extraversion and neuroticism during task-switching in young adults are consistent with previous findings and extend its implication to AD pathology possibly via differential regulation of neural activity throughout the lifetime, as discussed below.

### Personality, brain activity, and Aβ pathology

Accumulation of Aβ peptides is considered as an initiating event that precipitates downstream neuro/pathological changes in the process of AD^1,51^. Because of its significant impact in the pathogenesis of AD, there has been great interest in understanding the causal factors of Aβ accumulation during aging. One of the proposed mechanisms is increased neural activity and physiology early in life before plaque formation during senescence^52,53^. High topographic correspondence between increased metabolism in young people and increased Aβ deposition in older people has been consistently supported by both animal and human studies^18–20^. Incipient amyloid accumulation via neural activity may also lead to amyloidosis secondarily in other regions through prion-like spreading of the Aβ seed^51^. Therefore, an overall regulation of neural activity across the brain may be protective against brain Aβ deposition. Brain regions that accumulate brain Aβ include medial frontal and parietal cortices and anterior cingulate cortex that play a critical role in both cognitive and affective processes^49,54^. Dysfunction of these brain regions has been implicated in affective and psychiatric disorders such as depression and anxiety that are core personality characteristics associated with neuroticism^55^. Abnormal changes of these brain regions may relate to a dysfunction of the serotonergic system, which has been also suggested for amyloid precursor protein metabolism, or serotonin transporter polymorphisms^56,57^. Taken together, our findings provide evidence linking personality traits to Aβ pathology by showing differential lifelong regulations of brain activity in association with extraversion and neuroticism.

### Limitations and alternative explanations

Several other mechanisms might have mediated the observed relationship between personality traits and Aβ pathology. As personality traits such as extraversion have been significantly related to other personality traits such as openness to experience among older adults and conscientiousness in young subjects, it is possible that other lifelong variables may mediate the observed relationship between extraversion and lower Aβ pathology. Some genetic factors associated with personality may also explain its relationship with AD pathology via prevalent comorbidity associated with the genes^58^. More generally, personality may impact dementia risk and pathology by influencing health habits, number and quality of social relationships, cognitive activity, and reactions to stress over the lifespan, which could collectively have contributed to the observed relationship. In addition, because of the cross-sectional investigation of the current study, the observed association between extraversion and Aβ pathology in cognitively normal older adults may have reflected premorbid changes as other studies have suggested, although our prediction of fMRI data is against this possibility. Future studies are warranted to address these questions using a larger and longitudinal cohort and multivariate analysis approaches.

## Conclusion

The present study is the first to show that extraversion is associated with lower Aβ pathology among cognitively normal older adults and is associated with lower brain activity in regions including medial frontal cortex and anterior cingulate cortex in response to a controlled processing task in young adulthood. Our results provide evidence showing the neural basis underlying the link between extraversion and Aβ pathology that may develop throughout the lifespan. It is important to note that, although personality can be seen as a trait vs. state, it is potentially modifiable. In addition, while it is possible that life events, especially early stressful life events, may influence formation of personality type of individuals, one’s personality type also can modify the impact of stressful life events on health outcomes, affect lifestyle choices, and, more directly, impact AD pathology. Our results suggest that personality is an important factor that should be considered for conceptual and biological models of dementia risk and designing clinical trials. Considering its modifiability, personality can be considered for dementia risk reduction strategies that can start early in the lifespan.

## Acknowledgements

This research was supported by the National Institute on Aging (grant number R01AG026158). The author/s declare no competing financial interests.

## Supplementary materials

**Table S1.** Personality scores for five factor personality domains.

| **Personality** | **N** | **Minimum** | **Maximum** | **Mean** | **Std. Deviation** |
| --- | --- | --- | --- | --- | --- |
| **<u>All subjects</u>** |  |  |  |  |  |
| extraversion | 90 | 1.700 | 4.700 | 3.30333 | .766057 |
| agreeableness | 90 | 2.700 | 5.000 | 4.13877 | .632177 |
| conscientiousness | 90 | 2.000 | 5.000 | 3.88611 | .656882 |
| openness | 90 | 1.444 | 4.800 | 3.76694 | .658644 |
| neuroticism | 90 | 1.000 | 4.700 | 2.63086 | .896018 |
| **<u>Young subjects</u>** |  |  |  |  |  |
| extraversion | 33 | 1.700 | 4.700 | 3.39697 | .806414 |
| agreeableness | 33 | 2.700 | 5.000 | 4.10640 | .593685 |
| conscientiousness | 33 | 2.200 | 5.000 | 3.90909 | .655441 |
| openness | 33 | 1.444 | 4.700 | 3.81515 | .824279 |
| neuroticism | 33 | 1.300 | 4.700 | 2.90370 | 1.036344 |
| **<u>Older subjects</u>** |  |  |  |  |  |
| extraversion | 57 | 1.800 | 4.500 | 3.24912 | .743573 |
| agreeableness | 57 | 2.800 | 5.000 | 4.15750 | .657856 |
| conscientiousness | 57 | 2.000 | 5.000 | 3.87281 | .663165 |
| openness | 57 | 2.000 | 4.800 | 3.73904 | .546846 |
| neuroticism | 57 | 1.000 | 4.500 | 2.47290 | .770047 |

**Figure S1.**
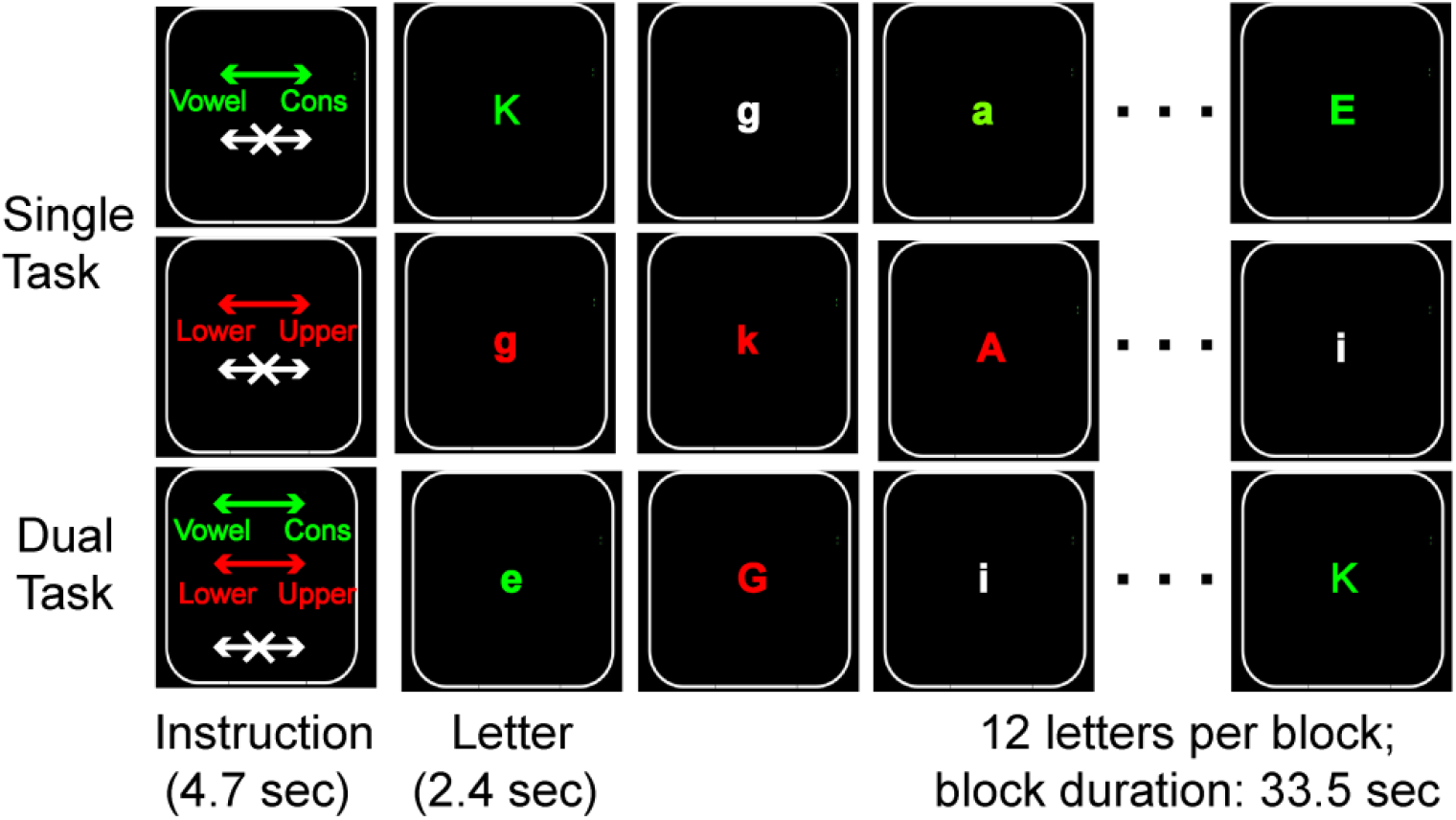
Schematic diagram of task-switching fMRI task.

